# Analysis of non-human primate models for evaluating prion disease therapeutic efficacy

**DOI:** 10.1101/2022.03.22.485331

**Authors:** Meredith A Mortberg, Eric Vallabh Minikel, Sonia M Vallabh

## Abstract

Prion disease is a fatal neurodegenerative disease caused by the conformational corruption of the prion protein (PrP), encoded by the prion protein gene (*PRNP*). While no disease-modifying therapy is currently available, genetic and pharmacological proofs of concept support development of therapies that lower PrP levels in the brain. In light of proposals for clinical testing of such drugs in presymptomatic individuals at risk for genetic prion disease, extensive nonclinical data are likely to be required, with extra attention paid to choice of animal models. Uniquely, the entire prion disease process can be faithfully modeled through transmission of human prions to non-human primates (NHPs), raising the question of whether NHP models should be used to assess therapeutic efficacy. Here we systematically aggregate data from N=527 prion-inoculated animals spanning six decades of research studies. Using this dataset, we assess prion strain, route of administration, endpoint, and passage number to characterize the relationship of tested models to currently prevalent human subtypes of prion disease. We analyze the incubation times observed across diverse models and perform power calculations to assess the practicability of testing prion disease therapeutic efficacy in NHPs. We find that while some models may theoretically be able to support therapeutic efficacy studies, pilot studies would be required to confirm incubation time and attack rate before pivotal studies could be designed, cumulatively requiring several years. The models with the shortest and most tightly distributed incubation times are those with smaller brains and weaker homology to humans. Our findings indicate that it would be challenging to conduct efficacy studies in NHPs in a paradigm that honors the potential advantages of NHPs over other available models, on a timeframe that would not risk unduly delaying patient access to promising drug candidates.

## Introduction

Prion disease is a rapidly fatal neurodegenerative disease of humans and other mammals. The pathogenic mechanism pivots on the conformational corruption of a host-encoded protein, the native prion protein or PrP, into a misfolded conformer, or prion, capable of corrupting other PrP molecules and killing neurons^1^. Uniquely, prion disease can arise in three ways. Sporadic cases (∼85%) appear to occur spontaneously, genetic cases (15%) trace to protein-coding variants in the prion protein gene, *PRNP* in humans^2^, and acquired cases (<1%), made famous by the kuru and variant CJD epidemics, can develop following iatrogenic exposure or consumption of prion-contaminated tissue^3^. The PrP dependence of all prion disease, regardless of etiology or even species, has long nominated the therapeutic hypothesis of PrP reduction^4^, and antisense oligonucleotides (ASOs) against the prion protein RNA now provide pharmacological proof of concept for this treatment strategy^5^. This progress motivates an assessment of available model systems in which to test PrP-lowering therapies.

The prion field benefits from unusually faithful animal models. Direct inoculation of animals with prion-infected brain homogenate induces the full prion disease process in which a clinically silent incubation period gives rise to characteristic symptoms, histopathology, and biochemical features, followed by terminal illness^6^. The inoculation paradigm has replicated across a range of mammalian systems, unified by key disease hallmarks and a fatal disease endpoint, but differing in time course and attack rate according to experimental parameters including inoculation route, prion strain^7^, species barrier^8^, and the PrP gene dosage of the host^9,10^. Over decades, prions have been bioassayed not only in a wide range of wild-type and transgenic rodent models, but in dozens of other mammals including cervids and non-human primates (NHPs)^11^. Despite this panoply of models, most studies have relied on intracerebral inoculation of mice with a well-characterized mouse-adapted prion strain, leveraging this system’s predictable time to disease^6^.

Given the rapid clinical progression of prion disease following symptom onset and ASO treatment data in mice suggesting an outsize benefit to early treatment^5^, it has been proposed that PrP-lowering agents could be tested clinically in presymptomatic individuals at known risk for genetic prion disease, with a goal of delaying or preventing onset^12^. Such a clinical path could involve the FDA’s Accelerated Approval program, in which a biomarker deemed “reasonably likely to predict clinical benefit” serves as the basis for provisional approval of a new drug. Because provisional approval could thereby precede direct observation of symptomatic benefit in humans, this strategy would likely demand unusually strong supporting data from animal models. The FDA’s “Animal Rule,” while designed for therapies unable to be tested in humans at all, and thus not directly applicable here, provides some insight into how regulators’ expectations for animal studies are adjusted when human efficacy studies are not feasible^13^.

The prospect of an unconventional clinical strategy draws special attention to the question of whether efficacy studies of such drugs in non-human primates (NHPs) would be feasible or advantageous. Unlike other non-transgenic models, a number of NHP species have been shown susceptible to human prion strains on direct passage from human tissue. Other theoretical advantages could include a *PRNP* sequence relatively closer to the human gene sequence, which might permit testing of a human DNA or RNA-targeting therapy in a non-transgenic animal, and a larger brain size better suited to simulating drug delivery to the human brain. The likelihood of meeting these interests would have to be balanced against concerns about achieving adequate power to reach a meaningful clinical endpoint in a large, onerous model; ensuring that variables such as prion strain and transmission route remain faithful to clinically relevant disease paradigms; and ensuring that the length of such a study would not unnecessarily delay access to human treatments.

In order to evaluate the prospects for efficacy studies in NHPs and assess how the above interests and tradeoffs might be balanced, we set out to exhaustively catalog and analyze reported NHP models. We began with a systematic literature search to identify published articles containing original data following prion-infected NHPs to disease endpoints. We then aggregated and manually curated a dataset of individual animal cohorts, and analyzed this dataset in order to i) determine how key experimental parameters in these models relate to prevalent forms of human prion disease, ii) analyze incubation times in these models and identify potential paradigms for efficacy studies, and iii) perform power calculations and assess the practicality and tradeoffs of various models.

## Methods

### Search strategy

To ensure a comprehensive and reproducible search, the following search strategy was adopted. All searches were conducted using the PubMed online database, between 2020-04-03 and 2020-12-22, with no date range imposed upon results. The search terms “non human primates,” “prions,” “inoculation,” “infected,” “Creutzfeldt-Jakob,” and “cynomolgus,” were used in combination. The initial results were supplemented by manual searches for the authors “Brown,” “Gajdusek,” “Marsh” and “Ono” to ensure that all work had been captured. See Supplemental Table 1 for search term and date details. Citations of relevant reviews^14,15^ were also screened. Titles and abstracts were reviewed for relevance, and only those containing primary data following prion-inoculated non-human primates to endpoint were included. Finally, manual follow-up was performed where reference lists in the identified reports suggested additional relevant titles.

Of 76 titles and abstracts reviewed (Figure 1), we excluded those lacking original NHP endpoint data not reported elsewhere (N=10), lacking any NHP data at all (N=9), lacking sufficient detail to determine outcomes for individual animals (N=3), or for which we were unable to obtain full text (N=2). We also excluded studies evaluating drug efficacy in primates (N=2): one^16^ was a conference abstract lacking experimental details, never subsequently published; the other^17^ reported treatment of prion-infected cynomolgus macaques with a novel small molecule compound, but provided no characterization of this animal model to justify that the two animals were sufficient to statistically power a conclusion regarding efficacy.

**Figure 1.**
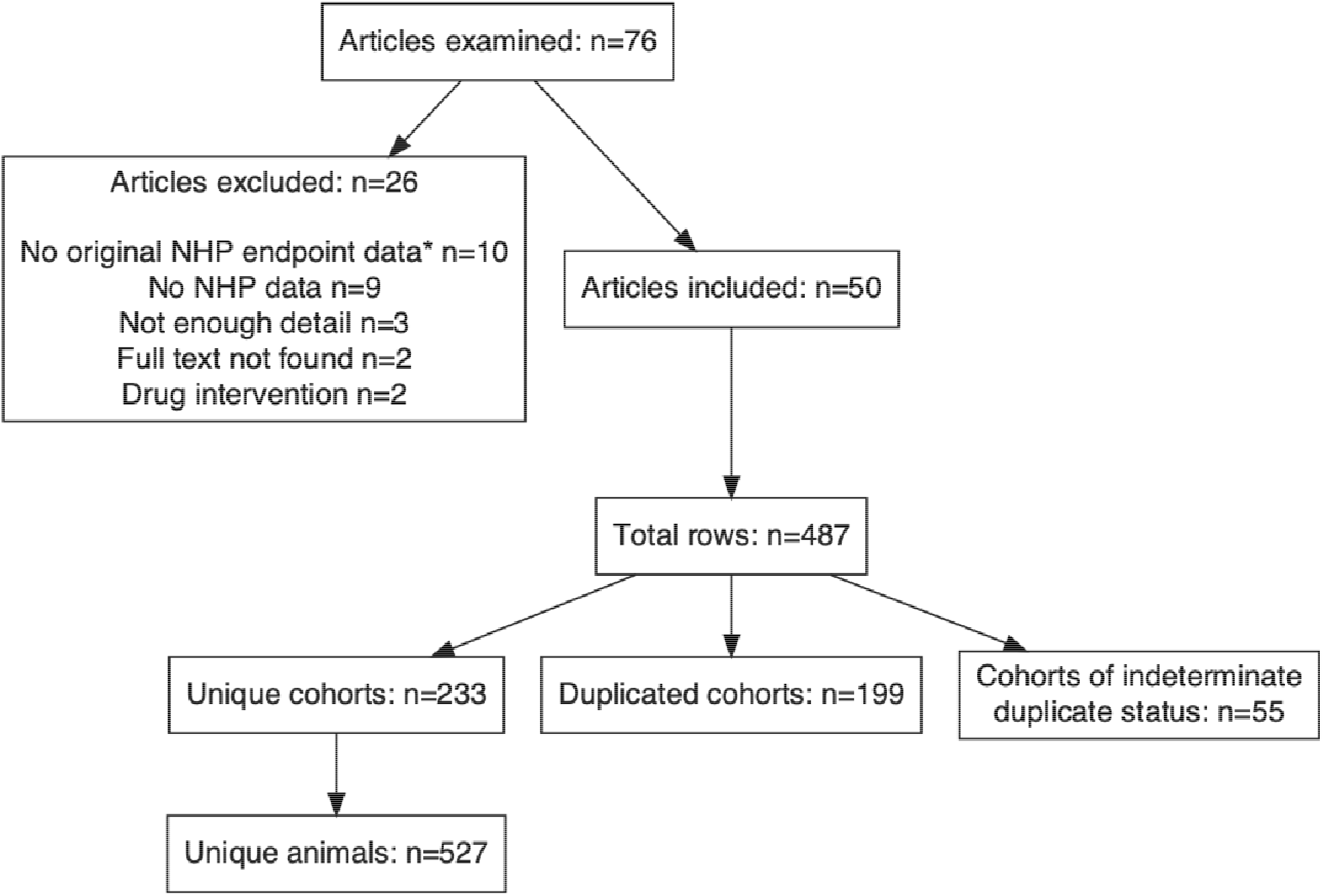
Schematic of the search strategy used to identify relevant articles.

Within the 50 remaining articles, one row was created for each unique report of an endpoint in a cohort of NHPs. Unique cohorts were defined as cohorts of animals of the same species, receiving the same prion inoculum by the same inoculation route. Many individual articles contained multiple unique cohorts, resulting in an initial count of N=487 rows when all reported cohorts were included. However, many NHP cohorts were the subject of multiple reports spanning years or decades, reflecting either multiple experimental endpoints (e.g. histology, symptom onset, and terminal illness) and/or published updates of experiments in progress. The rows were next manually de-duplicated with the goal of including any individual animal only once. This exercise identified both duplicated cohorts and cohorts for which insufficient details exist in the literature to determine whether or not they were elsewhere reported. Both were excluded, for a final count of N=233 unique cohorts comprising N=527 unique animals (Figure 1). The full dataset and species list are available as Supplemental Tables 2 and 3.

### Power calculations

Power calculation assumptions are enumerated under Results. For each scenario, we bootstrapped N=1,000 iterations and power was calculated as the percentage of those iterations in which a *P* value less than 0.05 was obtained. In each iteration, survival of untreated animals was sampled from a normal distribution with the reported mean and standard deviation, while survival of treated animals was sampled from a normal distribution with 1.5 times the reported mean and 1.5 times the reported standard deviation. For the “best case scenarios”, all animals were assumed to reach endpoint; for the “other scenarios”, a proportion (1-p) of animals were randomly censored, where p is the reported attack rate. Survival of treated and untreated animals was then compared using a two-sided log-rank test. For each iteration, the survival time of the longest-lived animal was also recorded. The expected study duration for each scenario was calculated as the average survival time of that longest-lived animal, across the 1,000 iterations.

### Homology analysis

Sequences for the *PRNP* gene in each species, from transcription start to stop including intronic and untranslated regions, were exported from UCSC Genome Browser, except for spider monkey, which was obtained from GenBank (PVHS01010010.1). Spider monkey and cynomolgus sequences, which are on the minus strand, were reverse complemented. The sequences were pairwise aligned to human *PRNP* using EMBOSS Needle^18^ with default parameters. Paired alignments were trimmed to remove any extraneous sequence context. Overall percent identity was calculated as the percent of human bases aligned as matches in the NHP species. The human gene was then tiled to generate every possible 20-mer, and if all 20 bases aligned as matches, the 20-mer was considered to have perfect identity.

### Statistical analysis and data availability

All analyses utilized custom scripts in R 4.0.4. Statistics in Figures 1, 2, and 3 are descriptive (N, mean, standard deviation, range) and are indicated in figure legends. Statistical tests and methods used in Table 1 are described under Power Calculations and Homology Analysis above. The curated dataset and source code sufficient to reproduce all analyses herein is available in a public git repository: https://github.com/ericminikel/nhp_models.

**Table 1.** Statistical power for efficacy studies in NHP models. ***(i): Best case scenarios from the literature***. *Studies from the prion NHP literature representing the most rapid model for the indicated combination of species, route of administration, and strain. Assuming 6 NHPs and a therapeutic that extends survival by 50%, estimates are given for mean time to endpoint, expected duration of study (time until the last animal reaches endpoint), and power*. ***(ii): Other scenarios with available data***. *Where available, other reported studies using the same species, strain, and inoculation route in at least N=3 NHPs are shown for comparison, along with estimates for mean time to endpoint, duration and power. *For sCJD i*.*c. in chimpanzees, the attack rate of 24/29 is limited to the animals included in the mean ± sd incubation time statistics provided by Brown et al; animals with longer incubation times up to 75 months are excluded*. ***(iii): Other considerations***. *For each paradigm, potential motivations for conducting an efficacy study in NHPs are evaluated. “Permitted” refers to whether the species is currently available for research in the United States. “Prevalent strain” refers to current clinical relevance of the prion strain. “Brain size” is calculated based on mass: human 1300g*^29^, *chimpanzee 387g*^30^, *cynomolgus 74g*^31^, *spider monkey 108g*^32^, *squirrel monkey 23g*^32^, *gray mouse lemur 2g*^33^. *“% identity” refers to percent sequence identify of each species’ full* PRNP *gene compared to the human* PRNP *gene. “% 20 bp runs identity” refers to the percent of twenty base-pair runs within each species’* PRNP *gene that are identical to the human* PRNP *gene*.

|  |  | (i)<br>best case scenario<br>from literature |  |  |  | (ii)<br>other scenarios with<br>available data |  |  |  |  | (iii)<br>other considerations |  |  |  |  |
| --- | --- | --- | --- | --- | --- | --- | --- | --- | --- | --- | --- | --- | --- | --- | --- |
| species | strain / RoA | study | survival time (mean±sd, months) | expected duration (months) | power | study | attack rate | survival time (mean±sd, months) | expected duration (months) | power | permitted | prevalent strain | brain size (% human) | overall % identity | % 20 bp runs identity |
| Chimpanzee | sCJD i.c. | <sup>24</sup> | 14.9±1.0 | 24 | 100% | <sup>25</sup> | 24/29* | 17±7* | 39 | 35% | n | y | 29% | 98.6% | 76.0% |
| Cynomolgus macaque | vCJD i.c. | <sup>26</sup> | 31.0±5.0 | 56 | 98% | | | | | | y | n | 5% | 97.5% | 33.8% |
| Spider monkey | sCJD i.c. | <sup>27</sup> | 23.5±0.8 | 37 | 100% | <sup>25</sup> | 30/31 | 31±8 | 61 | 73% | y | y | 8% | 90.0% | 16.9% |
| Squirrel monkey | sCJD i.c. | <sup>27</sup> | 24.6±3.7 | 44 | 98% | <sup>25</sup> | 196/211 | 25±5 | 47 | 87% | y | y | 2% | 89.0% | 15.0% |
| Gray mouse lemur | L-BSE i.c. | <sup>28</sup> | 20.6±1.6 | 34 | 100% | | | | | | y | n | 0.1% | 70.4% | 2.4% |

## Results

Our systematic literature search (Methods) identified N=50 publications reporting original data regarding prion disease endpoints in NHPs, totaling N=233 distinct animal cohorts and N=527 unambiguously unique individual animals (Figure 1). The temporal distribution of studies included in our analysis conformed to previous descriptions of two historical waves of primate research in the prion field^15^ (Figure 2A). The first wave, in the 1970s and 80s, corresponds to large scale inoculations performed largely at the National Institutes of Health, which have been deeply recounted elsewhere^19^. When divided by prion strain, kuru emerges as the major research interest of first wave, with more recent studies focused on transmission of bovine spongiform encephalopathy (BSE) and chronic wasting disease (CWD) (Figure 2B). Notably, considering that more than 99% of prion disease cases diagnosed today are sporadic or genetic, a minority of experimental primate inoculations have used a prion subtype currently affecting human patients; the kuru and BSE/vCJD epidemics are no longer major public health threats, and CWD has not been shown transmissible to humans.

**Figure 2.**
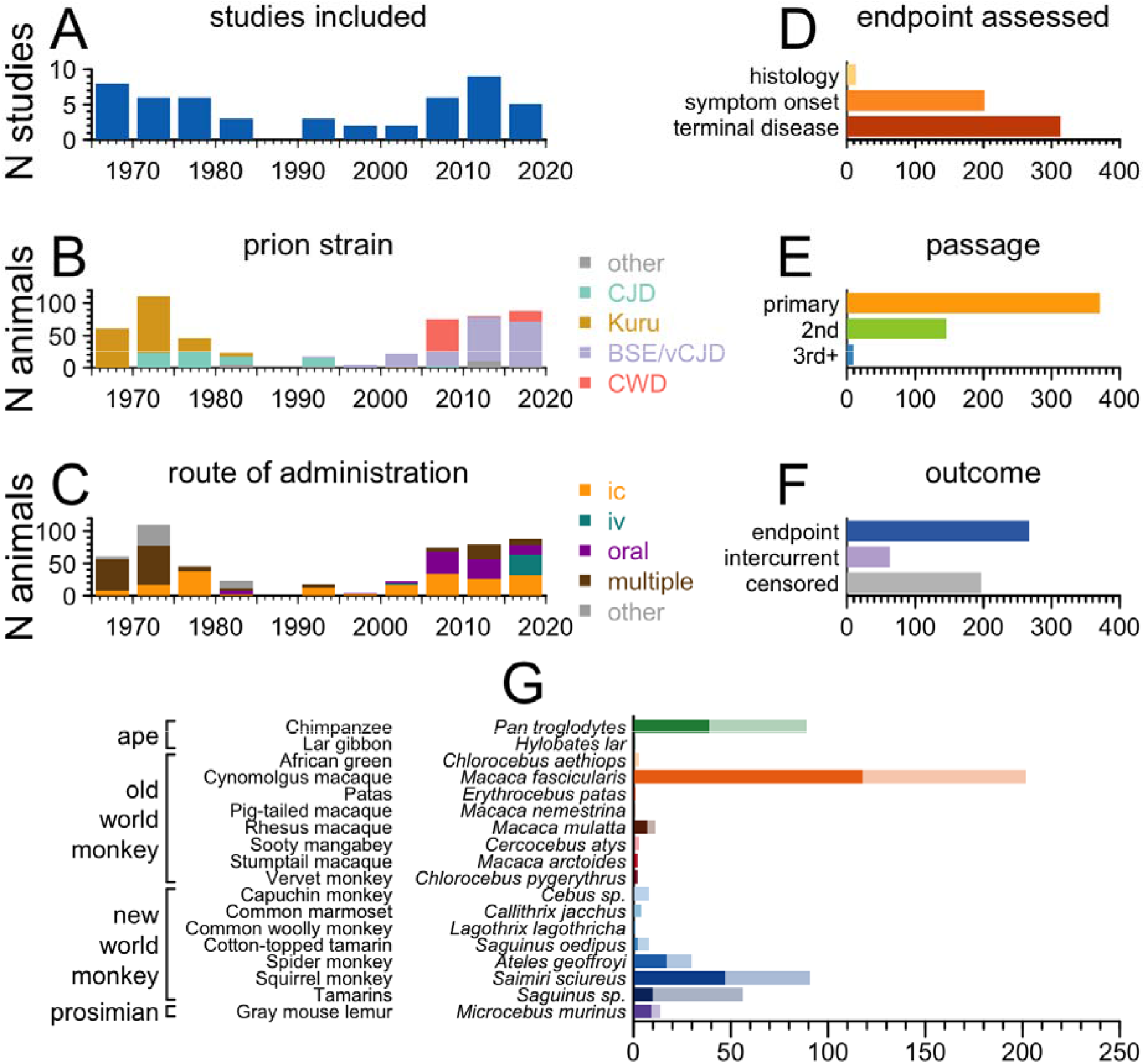
Overview of aggregated dataset of prion NHP experiments. Graphical summary of (A) studies included in our analysis by year; B) prion strains studied by year; C) routes of prion administration employed, by year; and D) study endpoints, by number of NHPs. E) Passage number of the prion inoculum used, by number of NHPs. Primary refers to direct human brain isolates. Second and third+ passage refer to inocula originating from human brain tissue, that have been inoculated into NHPs, then harvested and re-inoculated into subsequent NHPs. F) The number of NHPs reaching the study’s endpoint, lost to intercurrent illness, and censored at the time of study completion. G) The number of prion-inoculated NHPs reported in the prion literature, by species. Dark bars represent animals that reached endpoint, while light bars show animals that were lost to intercurrent illness or censored.

Today, intracerebral (IC) prion inoculation is considered the highest efficiency means of experimental transmission whether for primates^15^ or rodent models^6^. However, IC has not been the dominant inoculation method for primate studies (Figure 2C). When the parameters of transmission were still being explored, a wide range of techniques were tested and many animals were co-inoculated by more than one route. Meanwhile, oral inoculation is of special interest for BSE/vCJD and CWD, given that oral transmission led to zoonosis of BSE to humans^20^, and poses what is considered to be the greatest risk of zoonosis of CWD^21^. In addition, recent study of the intravenous (IV) method has been spurred by the discovery that vCJD has been transmitted via blood transfusion to four humans^22^. In total, 186/527 animals in our search were inoculated by the IC method alone.

Most animals reviewed were followed with the intention of observing a clinical endpoint of either symptoms or terminal disease (Figure 2D), following inoculation with a primary prion strain from a natural host, rather than a strain that had already undergone passage through non-human primates (Figure 2E.) Notably, however, given the length and difficulty of primate studies, roughly as many animals were either censored or lost to incurrent illness as were successfully followed to endpoint (260 vs. 267, Figure 2F).

Cynomolgus macaques were the most heavily represented primate species across studies. Chimpanzees, while well represented historically, are effectively no longer used for prion research following decisions by the NIH to phase out funding for chimpanzee research in 2013, and the U.S. Fish and Wildlife Service designation of all chimpanzees as endangered in 2015^23^. Squirrel monkeys therefore emerge as the second-most studied NHP species that remain amenable to research today (Figure 2G).

Broadly, NHP studies have sought to characterize prion transmission potential across diverse paradigms, spanning species, strains, and transmission routes (Figure 2). While it is clear that such studies take years, the time to endpoint varies both between and within experimental paradigms. 85% (197/233) of reported NHP cohorts have consisted of fewer than 4 animals, with 52% (121/233) of cohorts consisting of only 1 animal. If we limit our view to cohorts of at least N=3 animals, for which it is possible to estimate the distribution of survival times (see Figure 3 legend), then many combinations of species, strain and transmission route have been tested in only one experiment. For the paradigm that has been tested the greatest number of times – intracerebral inoculation of BSE into cynomolgus macaques – it is clear that these three variables alone do not standardize time to onset (Figure 3).

**Figure 3.**
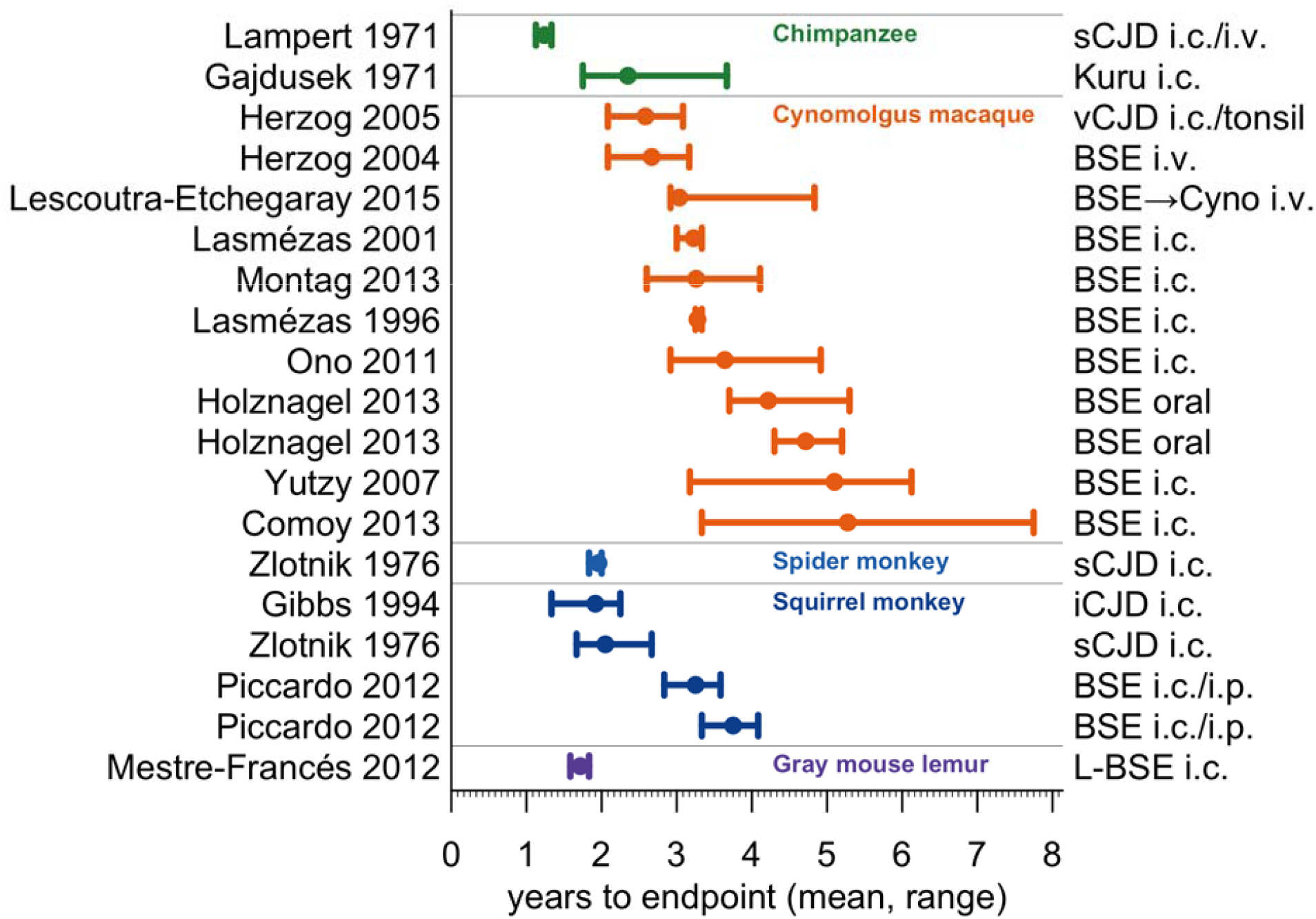
Duration of prion NHP studies. Cohorts for which it is possible to estimate the distribution of survival times were defined as those meeting all of the following criteria: i) containing at least N=3 animals total, ii) with at least N=3 reaching endpoint, iii) where all animals either reached endpoint or died of intercurrent illness, meaning none were censored at study termination, iv) where the endpoint studied was either terminal disease or symptom onset (as opposed to strictly histological outcomes), and v) for which the mean and standard deviation of survival time, or data sufficient to calculate such, were provided by the authors. The mean time to endpoint per cohort (dots), and range (bars), are shown alongside NHP species, strain and inoculation method.

For each of the five species represented in Figure 3, we selected the potentially most tractable combination of prion strain and route of administration for further analysis to determine the characteristics of a potential therapeutic efficacy study in each paradigm (Table 1). In order to calculate statistical power for such studies, we made the following assumptions:

1. Efficacy study of a therapeutic would require at least N=6 NHPs (N=3 of each sex, as Animal Rule Guidance recommends equal male and female representation).
2. Animals would be followed to terminal endpoint.
3. The therapeutic intervention would convey a 50% increase in survival time.
4. The outcome would be evaluated by log-rank survival test with a two-sided alpha=0.05 statistical threshold.
5. The study would last until the last animal reaches endpoint.

Based on these assumptions, we calculated the expected study duration and statistical power (1-β, probability of correctly rejecting the null hypothesis) for each paradigm under either i) the best-case scenario from the literature or ii) any other available reports, and tabulated these along with iii) other considerations for each model (Table 1). Other considerations (Table 1, Section iii) included whether use of the model is permitted for research, relevance of the prion strain, and brain size, as well as two metrics of homology to the human *PRNP* transcript. Because antisense oligonucleotide therapeutics, a modality currently in development for prion disease, are 20 base pairs long and are generally intolerant to single mismatches, we calculated not only overall percent identity, but also the percent of possible 20 base pair sequences that are 100% identical to the human sequence, a proxy for the probability of a drug designed for humans happening to match each species.

The best-case scenario for each paradigm would yield >80% power with a 2-5 year study duration, though it is possible that preliminary evidence of efficacy could be gleaned sooner. However, several caveats apply. First, as suggested by Figure 3, it is not clear that these combinations of species, strain, and inoculation route can be counted on to generate comparable results across studies. Indeed, where available, other reports in these paradigms suggest that attack rate may be lower, and/or incubation time longer, than observed in the “best case” report. The low, tightly distributed incubation time in the “best case” report for each paradigm might arise in part from luck, given the small number of NHPs in each cohort, and/or from properties of the exact brain sample inoculated, which may not still be available today.

In addition, not all motivations for performing an NHP study are satisfied by the paradigms highlighted in Table 2. Three out of five paradigms involve either a species (chimpanzees) no longer available, or a prion strain (vCJD or L-BSE) not responsible for many human prion disease cases today. Meanwhile increasing phylogenetic divergence from humans corresponds to steep drops in both NHP brain mass and *PRNP* sequence identify, particularly as measured in terms of the multi-base pair stretches of identity likely to be required to support targeting with a human-relevant genetically targeted therapy.

## Discussion

Transmission of human prions to NHPs is well established, and has been achieved independently by multiple laboratories dating back to the 1960s. Inocula representing multiple human prion strains have proven transmissible to multiple NHP species by multiple routes of inoculation. In this sense, NHP models of prion disease have been deeply explored. Many prion-inoculated cohorts of NHPs, however, consisted of only one or two animals per experimental condition, and/or exhibited incomplete attack rates or highly variable incubation times. Restricting our analysis to cohorts of three or more prion-inoculated NHPs that have developed terminal illness with a full attack rate reveals a more constrained universe of available models. Of these models, a handful have reached disease endpoint in two to three years on the mean, with standard deviations of only a few months. In particular, the best-case scenarios reported might suggest that a study in cynomolgus macaques, spider monkeys or squirrel monkeys could offer reasonable power to detect a therapeutic effect within a few years. For the latter two species, there is precedent for achieving this outcome with sCJD prions, corresponding to a dominant clinical subtype of human prion disease.

Nevertheless, our analysis highlights the challenges and limitations that such a study would face. While spider and squirrel monkeys appear generally susceptible to sporadic CJD prions, different studies have yielded incubation times of different magnitude and variability, potentially impacting study duration and statistical power (Table 1). This variability may derive from the fact most NHP experiments utilized primary passage of human prions (Figure 2E, Figure 3), with a distinct human brain isolate serving as inoculum in each study. Unlike in mice, where serial passage has given rise to well-characterized prion strains with typical incubation times and consistent terminal titers, distinct human brain isolates could reasonably be expected to differ in transmission-relevant properties such as prion titer and precise molecular subtype. The desire to ensure that a costly therapeutic efficacy study will not be wasted, should animal endpoints prove more variable than expected, might lead prudent sponsors to first conduct a pilot study to confirm incubation time for the exact prion isolate and inoculation procedure to be employed. Such a study would add years of up-front model development effort before a therapeutic efficacy study could begin, bringing the cumulative expected timeframe of a pilot study plus subsequent pivotal efficacy study to several years (Table 1). If such studies were gating for drug approval, they might unduly delay patient access to effective drugs.

Meanwhile, it is debatable whether spider or squirrel monkeys would honor all of the motivations for pursuing NHP studies to begin with. It is debatable whether they provide an advantage over non-NHP models in assessing drug brain distribution, as their brains are a small fraction of the mass of a human brain (8% and 2%, respectively) – smaller than those of sheep, goats, pigs, and large dogs^34^. Meanwhile, if efficacy studies need to be conducted with the actual human drug candidate rather than a surrogate compound^13^, the greater divergence of these New World monkeys from humans^35^ may pose an obstacle. In the context of a 20-base pair nucleic acid therapeutic, only 16.9% and 15.0% of the spider and squirrel monkey *PRNP* sequence, respectively, is composed of 20-meric runs of identity compared to the human gene. At this low level of identity, a drug targeting the human gene would be unlikely to show cross-reactivity by chance. Thus, cross-reactivity might only be achieved if it were prioritized in drug candidate selection, potentially compromising other drug parameters.

An alternative to the use of NHP prion infection models is to use separate models to address each question about a drug. Drug distribution in larger brains can be evaluated in uninfected NHPs or in other large animals. Efficacy of a human sequence-targeted therapeutic can be assessed in mice expressing human or chimeric *PRNP* genes. While the clarity and clinical utility of human prion disease sub-classifications are debated^2,36–38^, the existence of more than a dozen subtypes implies that any human prion strain chosen for an NHP study, even if transmitted directly from human inoculum, would still serve at best as a proxy for a subset of the patient population. Thus, work in a more facile model could serve to determine whether or not a given mechanism of action is strain-specific. To date, studies suggest that lowering levels of brain PrP, the substrate for formation and propagation of all prions, is effective across strains^5^.

The distribution of drug development activities across multiple more rapid and less costly models could allow a tighter feedback loop whereby insights from one experiment help inform a follow-on experiment, and allow a greater diversity of experimental parameters such as prion strain, drug dose, dosing regimen, and time of administration to be varied. By contrast, an efficacy study in an NHP model would require an intense investment of time and resources in a single experiment.

In summary, available data suggest that an NHP efficacy study of a prion disease therapeutic would be imaginable but daunting. The costs and benefits would need to be carefully weighed in light of both the drug type in question and the status of the drug development program to determine whether the scientific gains would outweigh the potential delay in advancing a therapeutic to human clinical trials.

## DISCLOSURES

EVM has received consulting fees from Deerfield Management and has received research support in the form of unrestricted charitable contributions from Ionis Pharmaceuticals. SMV has received speaking fees from Ultragenyx, Illumina, and Biogen, and has received research support in the form of unrestricted charitable contributions from Ionis Pharmaceuticals.

## ACKNOWLEDGMENTS

This study was funded by Prion Alliance, the Broad Institute (including direct philanthropic donations to Prions@Broad), the National Institutes of Health (R01 NS125255 to SMV), Ono Pharma Foundation, and an anonymous organization. We thank David Asher for his guidance in curation of the NHP prion literature.

## AUTHOR CONTRIBUTIONS

SMV conceived and designed the study, supervised the research, and wrote the manuscript. MAM conducted the literature search and curated the dataset. EVM performed homology analysis and power calculations. All authors analyzed the data, interpreted the results, edited and approved the final manuscript.

## Notes

https://github.com/ericminikel/nhp_models

## REFERENCES

1. Prusiner SB. Prions. PNAS. 1998 Nov 10;95(23):13363–13383. PMID: 9811807

2. Mead S, Lloyd S, Collinge J. Genetic Factors in Mammalian Prion Diseases. Annu Rev Genet. 2019 Dec 3;53:117–147. PMID: 31537104

3. Will RG. Acquired prion disease: iatrogenic CJD, variant CJD, kuru. Br Med Bull. 2003 Jun 1;66(1):255–265.

4. Lledo PM, Tremblay P, DeArmond SJ, Prusiner SB, Nicoll RA. Mice deficient for prion protein exhibit normal neuronal excitability and synaptic transmission in the hippocampus. Proc Natl Acad Sci U S A. 1996 Mar 19;93(6):2403–2407. PMCID: PMC39809

5. Minikel EV, Zhao HT, L. J, O’Moore J, Pitstick R, Graffam S, Carlson GA, Kavanaugh MP, Kriz J, Kim JB, Ma J, Wille H, Aiken J, McKenzie D, Doh-ura K, Beck M, O’Keefe R, Stathopoulos J, Caron T, Schreiber SL, Carroll JB, Kordasiewicz HB, Cabin DE, Vallabh SM. Prion protein lowering is a disease-modifying therapy across prion disease stages, strains and endpoints. Nucleic Acids Res. 2020 Aug 10;48(19):10615–10631. PMCID: PMC7641729

6. Watts JC, Prusiner SB. Mouse Models for Studying the Formation and Propagation of Prions. J Biol Chem. 2014 Jul 18;289(29):19841–19849. PMID: 24860095

7. Bartz JC. Prion Strain Diversity. Cold Spring Harb Perspect Med. 2016 Dec;6(12):a024349. PMCID: PMC5131755

8. Collinge J, Clarke AR. A general model of prion strains and their pathogenicity. Science. 2007 Nov 9;318(5852):930–936. PMID: 17991853

9. Büeler H, Aguzzi A, Sailer A, Greiner RA, Autenried P, Aguet M, Weissmann C. Mice devoid of PrP are resistant to scrapie. Cell. 1993 Jul 2;73(7):1339–1347. PMID: 8100741

10. Fischer M, Rülicke T, Raeber A, Sailer A, Moser M, Oesch B, Brandner S, Aguzzi A, Weissmann C. Prion protein (PrP) with amino-proximal deletions restoring susceptibility of PrP knockout mice to scrapie. EMBO J. 1996 Mar 15;15(6):1255–1264. PMCID: PMC450028

11. Brandner S, Jaunmuktane Z. Prion disease: experimental models and reality. Acta Neuropathol. 2017 Feb 1;133(2):197–222.

12. Vallabh SM, Minikel EV, Schreiber SL, Lander ES. Towards a treatment for genetic prion disease: trials and biomarkers. The Lancet Neurology. 2020 Apr 1;19(4):361–368.

13. U.S. Food and Drug Administration. Product Development Under the Animal Rule: Guidance for Industry [Internet]. 2015 Oct. Available from: https://www.fda.gov/regulatory-information/search-fda-guidance-documents/product-development-under-animal-rule

14. Trevitt CR, Collinge J. A systematic review of prion therapeutics in experimental models. Brain. 2006 Sep;129(Pt 9):2241–2265. PMID: 16816391

15. Krasemann S, Sikorska B, Liberski PP, Glatzel M. Non-human primates in prion research. Folia Neuropathol. 2012;50(1):57–67. PMID: 22505364

16. Amyx H, Salazar AM, Gajdusek DC, Gibbs CJ. Chemotherapeutic Trials in Experimental Slow Virus Diseases. Neurology. 1984. p. Suppl. 1 PP102.

17. Yamaguchi K, Kamatari YO, Ono F, Shibata H, Fuse T, Elhelaly AE, Fukuoka M, Kimura T, Hosokawa-Muto J, Ishikawa T, Tobiume M, Takeuchi Y, Matsuyama Y, Ishibashi D, Nishida N, Kuwata K. A designer molecular chaperone against transmissible spongiform encephalopathy slows disease progression in mice and macaques. Nat Biomed Eng. 2019 Mar;3(3):206–219. PMID: 30948810

18. Madeira F, Park YM, Lee J, Buso N, Gur T, Madhusoodanan N, Basutkar P, Tivey ARN, Potter SC, Finn RD, Lopez R. The EMBL-EBI search and sequence analysis tools APIs in 2019. Nucleic Acids Res. 2019 Jul 2;47(W1):W636–W641. PMCID: PMC6602479

19. Asher DM. Kuru: memories of the NIH years. Philos Trans R Soc Lond B Biol Sci. 2008 Nov 27;363(1510):3618–3625. PMCID: PMC2735519

20. Will RG, Ironside JW, Zeidler M, Cousens SN, Estibeiro K, Alperovitch A, Poser S, Pocchiari M, Hofman A, Smith PG. A new variant of Creutzfeldt-Jakob disease in the UK. Lancet. 1996 Apr 6;347(9006):921–925. PMID: 8598754

21. Hannaoui S, Schatzl HM, Gilch S. Chronic wasting disease: Emerging prions and their potential risk. PLoS Pathog. 2017 Nov 2;13(11):e1006619. PMCID: PMC5667731

22. Urwin PJM, Mackenzie JM, Llewelyn CA, Will RG, Hewitt PE. Creutzfeldt-Jakob disease and blood transfusion: updated results of the UK Transfusion Medicine Epidemiology Review Study. Vox Sang. 2016 May;110(4):310–316. PMID: 26709606

23. Grimm D. Chimps in waiting | Science. 2017 Jun 16;356(6343):1114–1117.

24. Lampert PW, Gajdusek DC, Gibbs CJ. Experimental spongiform encephalopathy (Creutzfeldt-Jakob disease) in chimpanzees. Electron microscopic studies. J Neuropathol Exp Neurol. 1971 Jan;30(1):20–32. PMID: 4925307

25. Brown P, Gibbs CJ, Rodgers-Johnson P, Asher DM, Sulima MP, Bacote A, Goldfarb LG, Gajdusek DC. Human spongiform encephalopathy: the National Institutes of Health series of 300 cases of experimentally transmitted disease. Ann Neurol. 1994 May;35(5):513–529. PMID: 8179297

26. Herzog C, Rivière J, Lescoutra-Etchegaray N, Charbonnier A, Leblanc V, Salès N, Deslys J-P, Lasmézas CI. PrPTSE distribution in a primate model of variant, sporadic, and iatrogenic Creutzfeldt-Jakob disease. J Virol. 2005 Nov;79(22):14339–14345. PMCID: PMC1280201

27. Zlotnik I, Grant DP, Dayan AD, Earl CJ, Illis LS, Weller RO. Further observations on the experimental transmission of Creutzfeldt-Jakob disease from man to squirrel and spider monkeys. Neuropathology and Applied Neurobiology. 1976;2(2):125–130.

28. Mestre-Francés N, Nicot S, Rouland S, Biacabe A-G, Quadrio I, Perret-Liaudet A, Baron T, Verdier J-M. Oral transmission of L-type bovine spongiform encephalopathy in primate model. Emerg Infect Dis. 2012 Jan;18(1):142–145. PMCID: PMC3310119

29. Svennerholm L, Boström K, Jungbjer B. Changes in weight and compositions of major membrane components of human brain during the span of adult human life of Swedes. Acta Neuropathol. 1997 Oct;94(4):345–352. PMID: 9341935

30. Herndon JG, Tigges J, Anderson DC, Klumpp SA, McClure HM. Brain weight throughout the life span of the chimpanzee. J Comp Neurol. 1999 Jul 12;409(4):567–572. PMID: 10376740

31. Pardo ID, Garman RH, Weber K, Bobrowski WF, Hardisty JF, Morton D. Technical guide for nervous system sampling of the cynomolgus monkey for general toxicity studies. Toxicol Pathol. 2012 Jun;40(4):624–636. PMID: 22317925

32. Hartwig W, Rosenberger AL, Norconk MA, Owl MY. Relative brain size, gut size, and evolution in New World monkeys. Anat Rec (Hoboken). 2011 Dec;294(12):2207–2221. PMID: 22042631

33. MacLean EL, Barrickman NL, Johnson EM, Wall CE. Sociality, ecology, and relative brain size in lemurs. J Hum Evol. 2009 May;56(5):471–478. PMID: 19410273

34. Crile G, Quiring Daniel P. A Record of the Body Weight and Certain Organ and Gland Weights of 3690 Animals. Ohio Journal of Science. 1940 Sep;40(5):219–259.

35. Perelman P, Johnson WE, Roos C, Seuánez HN, Horvath JE, Moreira MAM, Kessing B, Pontius J, Roelke M, Rumpler Y, Schneider MPC, Silva A, O’Brien SJ, Pecon-Slattery J. A molecular phylogeny of living primates. PLoS Genet. 2011 Mar;7(3):e1001342. PMCID: PMC3060065

36. Collins SJ, Sanchez-Juan P, Masters CL, Klug GM, van Duijn C, Poleggi A, Pocchiari M, Almonti S, Cuadrado-Corrales N, de Pedro-Cuesta J, Budka H, Gelpi E, Glatzel M, Tolnay M, Hewer E, Zerr I, Heinemann U, Kretszchmar HA, Jansen GH, Olsen E, Mitrova E, Alpérovitch A, Brandel J-P, Mackenzie J, Murray K, Will RG. Determinants of diagnostic investigation sensitivities across the clinical spectrum of sporadic Creutzfeldt-Jakob disease. Brain. 2006 Sep;129(Pt 9):2278–2287. PMID: 16816392

37. Parchi P, Strammiello R, Notari S, Giese A, Langeveld JPM, Ladogana A, Zerr I, Roncaroli F, Cras P, Ghetti B, Pocchiari M, Kretzschmar H, Capellari S. Incidence and spectrum of sporadic Creutzfeldt–Jakob disease variants with mixed phenotype and co-occurrence of PrPSc types: an updated classification. Acta Neuropathol. 2009 Nov 1;118(5):659–671.

38. Takada LT, Kim M-O, Cleveland RW, Wong K, Forner SA, Gala II, Fong JC, Geschwind MD. Genetic prion disease: Experience of a rapidly progressive dementia center in the United States and a review of the literature. Am J Med Genet B Neuropsychiatr Genet. 2017 Jan;174(1):36–69. PMCID: PMC7207989

